# Reconciling the spectrum of RORγt^+^ antigen-presenting cells

**DOI:** 10.1101/2023.11.01.565227

**Authors:** Tyler Park, Christina Leslie, Alexander Y. Rudensky, Chrysothemis C. Brown

## Abstract

Three recent contemporaneously published papers^1–3^ showed that antigen-presenting cells (APC) expressing the nuclear receptor RORγt instruct peripheral regulatory T (pTreg) cell differentiation and establish tolerance to the gut microbiota. These studies identified a spectrum of RORγt^+^ APCs, distinct from dendritic cells, that included type 3 innate lymphoid cells (ILC3s) as well as novel APC types. Given the discordant conclusions as to the nature of the pTreg-inducing APC and the divergent nomenclature used in these three studies, there is a clear need to reconcile these analyses and the identity of the cell types described. Our reanalysis of the single cell RNA-seq data from these studies revealed the presence of four distinct subsets of non-ILC3 RORγt^+^ APCs, present in all three datasets reported, and confirmed expression of Itgb8, the critical factor for RORγt^+^ APC mediated pTreg induction, in cells synonymous with the non-Aire expressing Thetis cell subset, TC IV.

---

Three recent contemporaneously published papers^1–3^ showed that antigen-presenting cells (APC) expressing the nuclear receptor RORγt instruct peripheral regulatory T (pTreg) cell differentiation and establish tolerance to the gut microbiota. These studies identified a spectrum of RORγt^+^ APCs, distinct from dendritic cells, that included type 3 innate lymphoid cells (ILC3s) as well as novel APC types. Given the discordant conclusions as to the nature of the pTreg-inducing APC and the divergent nomenclature used in these three studies, there is a clear need to reconcile these analyses and the identity of the cell types described. In the study by Lyu et al.^3^, the authors found that RORγt^+^ APCs included lymphoid tissue inducer (LTi)-like cells (ILC3s) as well as two clusters of cells termed RORγt^+^ extra-thymic Aire-expressing cells (eTACs). In the study by Kedmi et al.^2^, aside from LTi-like cells, RORγt^+^ APCs encompassed a population of Aire-expressing cells termed Janus cells (JC) based on the earlier description of RORγt^+^Aire^+^ JCs by Wang et al.^4^. In addition to LTi cells, Akagbosu et al. identified 4 distinct subsets of cells, termed Thetis cells (TC I-IV) that included two subsets of Aire-expressing cells (TC I and TC III)^1^. Lyu et al. ascribed pTreg cell inducing capacity and maintenance of intestinal tolerance exclusively to ILC3. To the contrary, Akagbosu et al. implicated a specific subset of TCs – Aire-negative TC IV – while excluding a contribution from ILC3s. Finally, Kedmi et al. suggested a central role for RORγt^+^ APCs, either ILC3 or other RORγt^+^ cells such as Aire-expressing JCs, in these processes. While differences exist between the reporters used for identification of the cells (Aire-GFP vs RORγt-GFP vs RORc-Venus), the age of the mice analyzed, and the lymph node of origin – factors that affect the relative abundance of each of the cell types – sufficient commonality across the datasets allowed for their meaningful comparison. Thus, to reconcile these three studies we undertook a re-analysis of their publicly available datasets.

To determine the relationships between the cells described, we analyzed each dataset individually and then integrated the datasets for direct comparison. We first turned to the scRNA-seq analysis of RORγt(GFP)^+^ cells isolated from mesenteric lymph nodes (mLN) of adult mice by Lyu et al. Within this dataset, RORγt^+^MHCII^+^ cells spanned a cluster of LTi-like cells and two additional clusters of cells denoted as ‘RORγt eTAC I’ and ‘RORgt eTAC II’ (**Fig. 1a**). These ILC3 and non-ILC3 subsets were distinguished by differential Rora and Cxcr6 expression (**Fig. 1b**), consistent with the data from Akagbosu et al. Analysis of expression of TC I-IV signature genes, defined by Akagbosu et al., showed that RORγt eTAC I expressed signature genes for Aire^+^ TC I (*Nrxn1, Gal*), and included a small number of cells, separated in UMAP space, expressing signature TC II genes (*Col17a1, Hk2*). The sparse RORγt^+^ eTAC II cluster compromised a heterogeneous group of cells expressing TC III (*Dnase1l3, Nlrc5*) or TC IV marker genes (*Itgb8* and *Ccl22*) (**Fig. 1c**), suggesting that the low cell numbers may have obscured the identification of discrete subsets of cells through conventional clustering methods. The expression of Itgb8, although not noted by Lyu et al., is significant given that a requirement for Itgb8 expression by RORγt^+^ APCs for pTreg differentiation was established by two other studies^1,2^.

**Figure 1.**
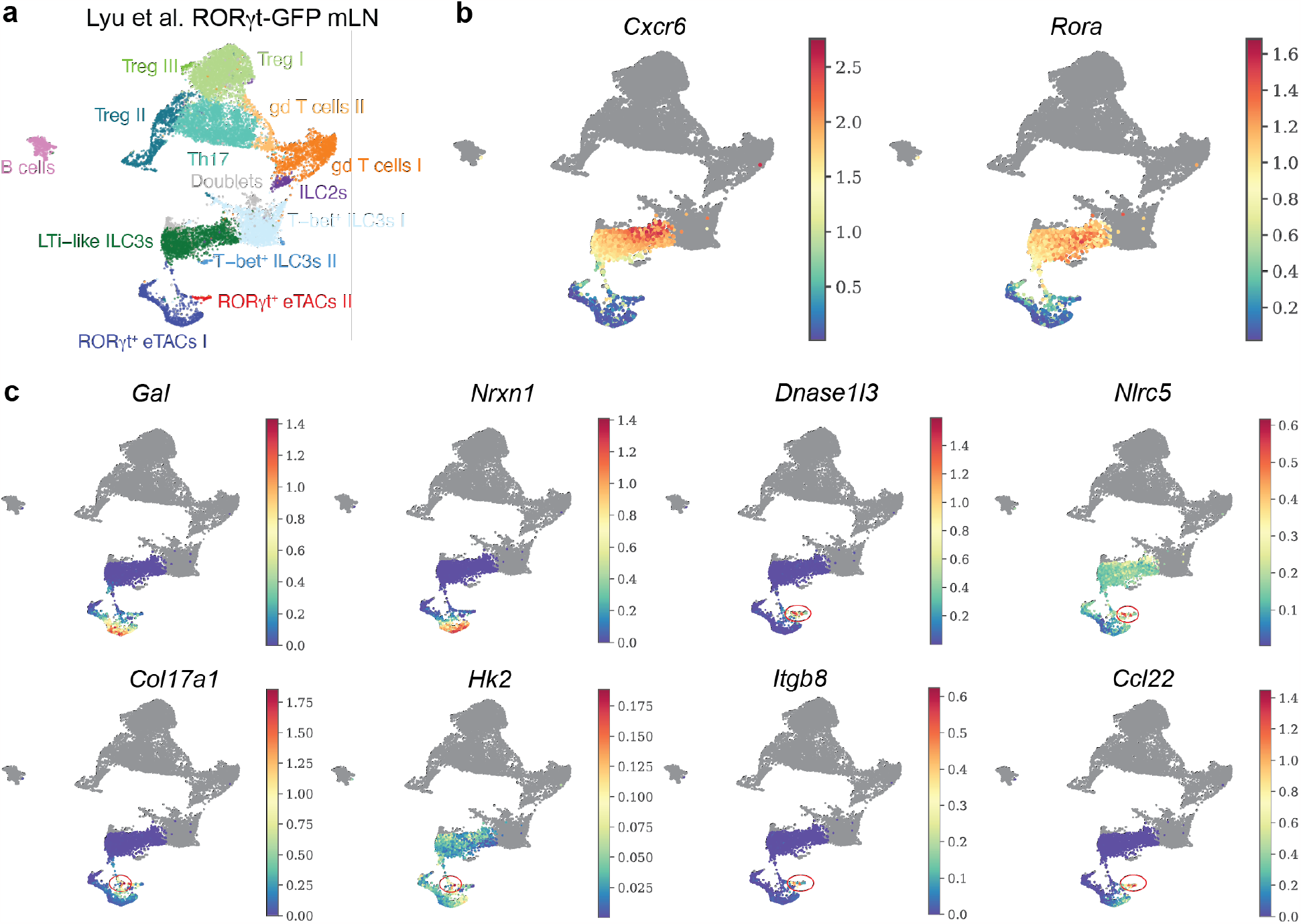
Reanalysis of RORγt-GFP sequencing data made publicly available by Lyu et al. **a-b**. UMAP visualization of scRNA-seq analysis from RORγt-GFP^+^ mesenteric lymph node (mLN) cells from Lyu et al., colored (a) by original cell type annotation or (b) imputed expression of ILC3 genes, CXCR6 and RORα . **c**, UMAP visualization of RORγt-GFP^+^ mLN cells from Lyu et al. colored by imputed expression of TC subset signature genes.

In the Kedmi et al. study, RORγt^+^ APCs included LTi-like ILC3s and non-ILC3 cells (JC) encompassing three distinct clusters (JC1-3). Of note, JCs were identified through scRNA-seq analysis of GFP^+^ cells isolated from a BAC-transgenic Aire-GFP reporter mouse^5^. The reported analysis combined cells from two separate datasets: one previously published by Wang et al.^4^ and an independently generated dataset. Considering the combined data by necessity underwent batch correction which has the potential to blunt both technical and biological variation in gene expression, we analyzed each dataset individually. Focusing on the most recent BAC Aire-GFP^+^ cell dataset (GSE200148), we identified 5 clusters of RORγt -expressing MHCII^+^ cells (R1-R5) (**Fig. 2a,b**). Cluster R5 cells were characterized by lower transcript numbers, a higher fraction of mitochondrial genes and increased expression of the apoptotic marker gene *Malat1*, and were therefore excluded from downstream analysis. Amongst the remaining RORγt^+^MHCII^+^ clusters, analysis of ILC3 signature genes revealed one cluster of CXCR6^+^ LTi cells (R1), and three clusters of CXCR6^−^ non-ILC3 cells (R2-R4) (**Fig. 2c**). Cluster R2 comprised proliferating (Ki67^+^) cells, cluster R3 expressed TC I signature genes, and cluster R4 contained a heterogeneous group of cells expressing marker genes of either TC II, III, or IV (**Fig. 2c**,**d**). Overlaying these clusters with the signatures for JC1-3 subsets defined by Kedmi et al., we found that the cells within R2 (TC-I like cluster) were distinguished by the JC1 transcriptional signature (**Fig. 2e**), while the JC2 signature was enriched in the heterogeneous R3 cluster. The JC3 signature mapped to the R5 cluster containing low QC cells (**Fig. 2e**). We next analyzed the Aire-GFP dataset from the same study that was first reported by Wang et al. and similarly identified four clusters of RORγt-expressing cells (**Fig. 3a,b**). Cluster R1 represented CXCR6^+^ LTi cells, cluster R2 contained proliferating (Ki67^+^) cells, R3 expressed signature TC I genes and cluster 4 contained a mixture of cells with TC II-IV signature genes, again separated in UMAP space suggesting potentially distinct cell types (**Fig. 3c,d**). Overlay of JC1, 2, or 3 signatures confirmed that JC1 corresponded to TC I (**Fig. 3e**). As before, the JC2 signature was enriched in the mixed cluster R4 (**Fig. 3e**) while JC3 enriched cells were not identified in this dataset.

**Figure 2.**
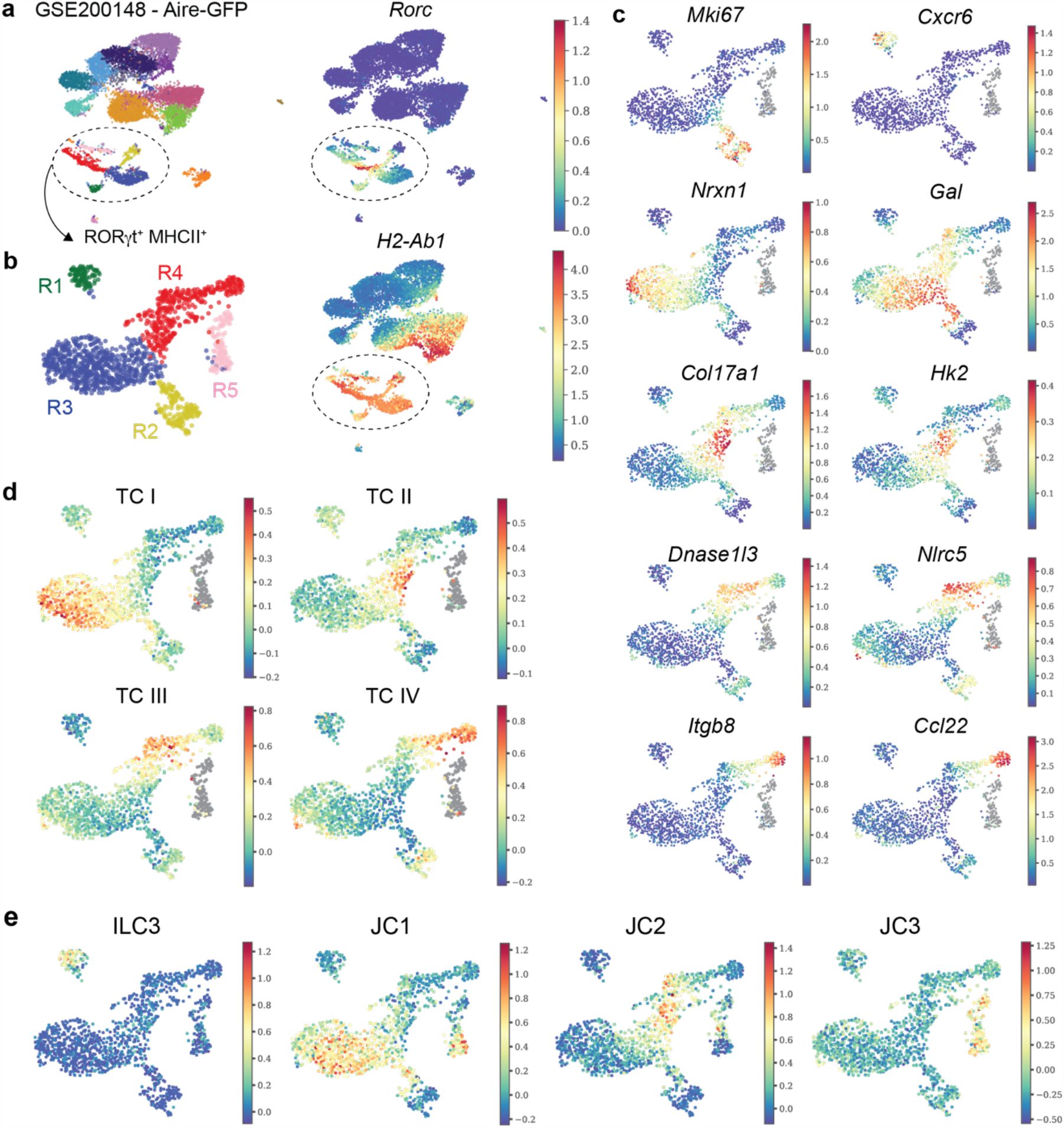
Reanalysis of Aire-GFP sequencing data made publicly available by Kedmi et al. **a**. UMAP visualization of scRNA-seq analysis from Aire-GFP^+^ lymph node cells from Kedmi et al., colored by cluster annotation or imputed expression of Rorc and H2-Ab1. **b**. UMAP visualization of RORγt^+^MHCII^+^ clusters. **c**-**e**, UMAP visualization of RORγt^+^MHCII^+^ cells colored by imputed expression of select TC subset genes (c), unimputed TC subset signature score (d), or unimputed JC subset signature score (e).

**Figure 3.**
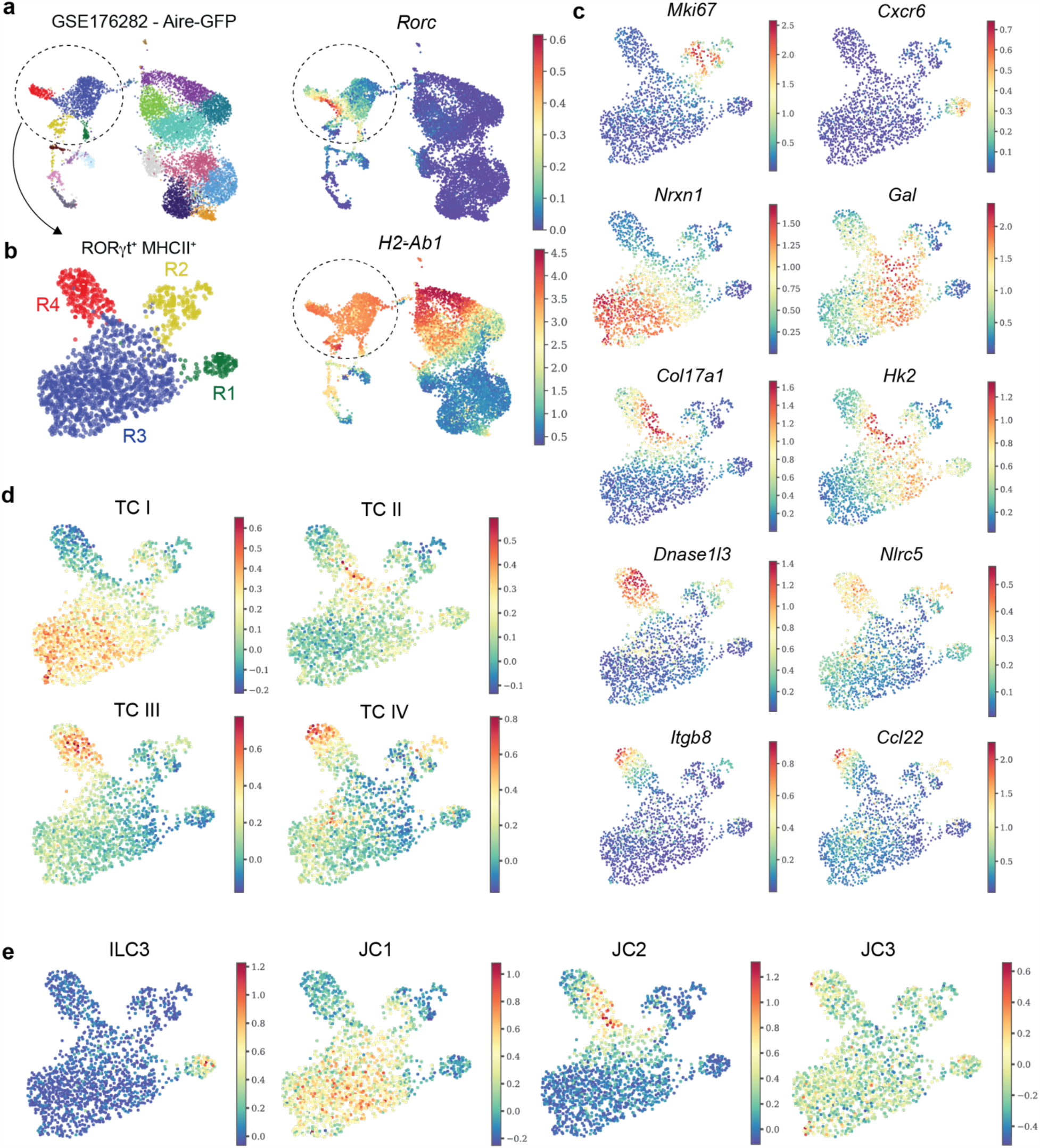
Reanalysis of Aire-GFP sequencing data made publicly available by Wang et al. **a**. UMAP visualization of scRNA-seq analysis from Aire-GFP^+^ lymph node cells from Wang et al., colored by cluster annotation or expression of Rorc and H2-Ab1. **b**. UMAP visualization of RORγt^+^MHCII^+^ clusters. **c**-**e**, UMAP visualization of RORγt^+^MHCII^+^ cells colored by imputed expression of select TC subset genes (c), unimputed TC subset signature score (d), or unimputed JC subset signature score (e).

Thus, the common findings upon re-analysis of all three datasets were the presence of Aire-expressing RORγt^+^ cells with features of TC I and clusters of cells with mixed TC II-IV marker genes. A considerable challenge to identifying distinct cell types in a heterogenous mix of cells is the sparsity of TCs in datasets generated using 4w or adult mice^2,3^, where their numbers are ∼5 fold lower than their peak abundance in mice at 2 weeks of age^1^. Therefore, to achieve sufficient power to resolve the identity of the cell types identified in the three studies, we pooled all RORγt^+^MHCII^+^ cells across the four datasets and performed single cell integration, using established methods.

Visualization of the integrated data (**Fig. 4a**) demonstrated that cells identified as LTi/ILC3 cells in all three studies mixed together, confirming that the integration method has the ability to accurately map the same type of cells with similar transcriptional program. Consistent with their expression of TC I signature genes, the majority of cells defined as RORγt^+^ eTAC I (Lyu et al.) and JC1 (Kedmi et al.) mapped with TC I cells. Notably, cells from the Lyu et al. RORγt^+^ eTAC II cluster segregated with either TC III or IV cells, in accordance with their expression of corresponding gene signatures. Similarly, the R4 cluster of cells identified in the two datasets from Kedmi et al. mapped to corresponding TC II, III or IV clusters.

**Figure 4.**
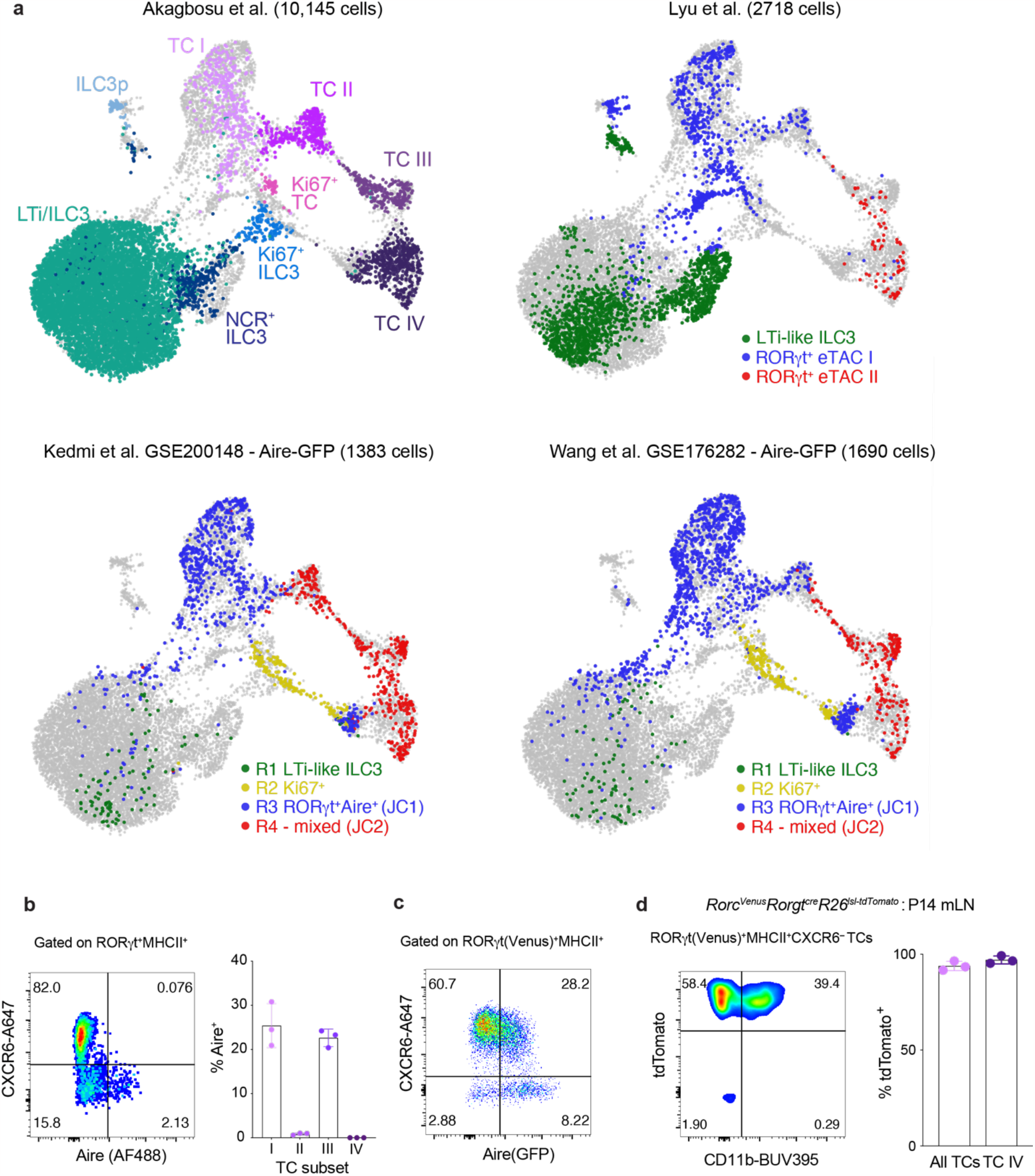
Integrated analysis of RORγt^+^MHCII^+^ scRNA-seq data. **a**. UMAP visualization of RORγt^+^MHCII^+^ cells from Akagbosu et al, Kedmi et al. and Lyu et al. colored by original cell type annotation (Akagbosu et al. and Lyu et al.) or cluster annotation from Fig. 2b and 3b (Kedmi et al.). b. Representative flow cytometry analysis of intraceullular Aire protein expression in CXCR6^+^MHCII^+^ILC3 and CXCR6^−^TCs isolated from mLN at P14, and summary bar graphs. Each dot represents an individual mouse (*n* = 3). **c**, Representative flow cytometry analysis of Aire-GFP expression in CXCR6^+^MHCII^+^ILC3 and CXCR6– TCs isolated from mLN of *Rorc*^Venus^Aire^GFP^ mice at P14. **d**, Flow cytometry of Lin^−^RORγt(Venus)^+^CXCR6^−^MHCII^+^ TCs from mLN of *Rorgt*^*Cre*^*R26*^*lsl-tdtomato*^*Rorc*^*Venus-CreERT2*^ mice and summary graph of frequency of tdTomato^+^ cells amongst all TCs and CD11b^+^ TC IV (*n* = 3 mice). Data in **b** and **c** are representative of >5 independent experiments. Data in **c** are representative of 2 independent experiments.

Together, these analyses resolve the apparent discrepancies between these three studies, which at first glance appear to describe different populations of RORγt^+^APCs, by confirming that the non-ILC3 RORγt^+^MHCII^+^ cell population comprises 4 distinct cell types. Clearly the timing of sampling is critical for determining the phenotype of these cells due to their enrichment in early life and lesser abundance in adult mice^1^. The numerically dominant non-ILC3 RORγt^+^ type in the latter appears to be Aire^+^RORγt^+^ cells which represent TC I (synonymous with RORγt^+^ eTAC I or JC1). In addition to this Aire-expressing cell type, there are 3 additional sub-types of non-ILC3 RORγt^+^ APCs, present in all 3 datasets albeit rare in adult mice, representing TC II-IV. Finally, JC2 or RORγt^+^ eTAC II signatures relate to mixed populations of TC II-IV cells rather than defining a discrete cell type or state. In our opinion, the phenotypic characterization combined with unifying nomenclature, which allows subdivision of novel RORγt^+^ APCs into 4 distinct cell types and, importantly, delineates Aire^−^ TC IV and TC II as well as transcriptionally divergent Aire-expressing TC I and TC III cell types, are critical for future elucidation of the developmental pathways and functions for these distinct cell types. Of note, only two of these cell types meet the definition of ‘extrathymic Aire expressing cells (eTAC)’ based on significant transcript and Aire protein expression. As noted by previous studies, expression of BAC-Aire-GFP is not synonymous with Aire protein expression^1,6^. To further investigate the expression of Aire in RORγt^+^ APCs and accuracy of RORγt^+^ eTACs as a term for non-ILC3 RORγt^+^ APCs, we compared expression of Aire protein in wild-type RORγt^+^MHCII^+^ cell types vs expression of BAC-Aire-GFP. Analysis of Aire protein expression confirmed that Aire is exclusively expressed by two subsets of CXCR6^−^RORγt^+^MHCII^+^ cells – TC I and III (**Fig. 4b**), as previously demonstrated^1^. In contrast, analysis of Aire-GFP expression revealed that over 30% of CXCR6^+^RORγt^+^MHCII^+^ LTi/ILC3s expressed the Aire-GFP transgene, indicating discrepant expression of this reporter and endogenous Aire protein (**Fig. 4c**). Overall, these analyses demonstrate that the term ‘RORγt^+^ eTAC’ is only applicable to two of the four subsets of non-ILC3 RORγt^+^ APCs.

Finally, it has recently been suggested by Abramson et al. that TC IV i) may not express the RORγt isoform and ii) that their existence has not been independently confirmed by other studies^7^. However, we now show that Itgb8-expressing TC IV cells were captured and profiled in the scRNA-seq analysis by Lyu et al, in which cells were sorted on the basis of GFP expression driven by the RORγt promoter. This is in full agreement with our previously reported highly efficient fate mapping of TCs by Rorgt^cre^ (95%) including TC IV (96-99%; **Fig. 4d**). As noted by Abramson et al in a recent discussion of these three studies, “new tools to specifically and robustly target each novel RORγt^+^ APC subset are needed to clarify their ontogeny, lineage relationships and functional potential”. Our reconciliation of the 3 datasets, and clarification on the existence of four discrete non-ILC3 cell types in all studies aids in the development of these tools, allowing functional dissection of each subset.

## Methods

### Single-cell RNA-seq computational analysis

For the scRNA-seq dataset from Lyu et al. (GSE184175), cells annotated as RORγt^+^MHCII^+^ cell types (LTi-like ILC3, RORγt eTAC I and RORγt eTAC II) in the original study were selected for downstream analysis. For the scRNA-seq dataset from Akagbosu et al. (GSE174405), cells annotated as TC and ILC3 cell types in the original study were selected for analysis. For the scRNA-seq datasets from Kedmi et al. (GSE200148 and GSE176282), the samples containing cells from Aire-GFP lymph nodes were selected for analysis. For each dataset from Kedmi et al., the count matrix was filtered based on the number of transcripts (>1,000 and <30,000), the number of detected genes (>600 and <5,000), and the fraction of mitochondrial transcripts (<10%). The filtered count matrix was library-size normalized, log-transformed (‘log-normalized’ expression values) and then centered and scaled (‘scaled’ expression values) using Seurat v4.3.0. Principal component analysis (PCA) was performed on the scaled data (npcs=50). PhenoGraph clustering was performed using the first 30 principal components (PCs) with 30 nearest neighbors. Cell clustering was visualized using UMAP, computed from the nearest neighbor graph built by PhenoGraph.

For visualization of gene expression, MAGIC imputation was applied to the log-normalized expression values to further de-noise and recover missing values. Imputed gene expression values were only used for data visualization of individual genes on UMAP overlays. TC and JC signature scores were computed using ‘AddModuleScore’ function with default parameters from Seurat. The top 130 differentially expressed genes in each of the TC subsets were used as TC signature genes. Differentially expressed genes reported in Kedmi et al. (Extended Data Fig. 7d) were used as signature genes for ILC3 and JC subsets.

### Integrating the single-cell RNA-seq datasets

The four scRNA-seq datasets were integrated using Seurat’s anchor-based integration method. First, 5,000 genes were first selected based on their repeated variability across datasets to identify anchors using canonical correlation analysis. The integration of the datasets was then performed using ‘IntegrateData’ function. Next, the expression values of genes in the integrated dataset were scaled and used for PCA. A UMAP embedding of the integrated dataset was computed based on the top 30 principal components and used for visualization of the cells.

### Mice

*Rorc*^*Venus-T2A-creERT2*^ and Adig(Aire^GFP^) mice have been previously described^1,5^. *Rorgt*^*cre*^, *R26*^*lsl-tdTomato*^, C57Bl/6 mice were purchased from Jackson Laboratories. Generation and treatments of mice were performed under protocol 21-05-007, approved by the Sloan Kettering Institute (SKI) Institutional Animal Care and Use Committee.

### Tissue processing

Mice were euthanized by CO_2_ inhalation. Organs were harvested and processed as follows. Lymphoid organs were digested in collagenase in RPMI1640 supplemented with 5% fetal calf serum, 1% L-glutamine, 1% penicillin–streptomycin, 10 mM HEPES, 1 mg/ml collagenase A (Sigma, 11088793001) and 1U/mL DNase I (Sigma, 10104159001) for 45 min at 37°C, 250 rpm. Digested samples were filtered through 100-μm strainers and centrifuged to remove collagenase solution.

### Flow cytometry

For flow cytometric analysis, dead cells were excluded either by staining with LIVE/DEAD Zombie NIR in PBS for 10 minutes at 4°C, prior to cell-surface staining. Cells were then incubated with anti-CD16/32 in staining buffer (2% FBS, 0.1% Na azide, in PBS) for 10 minutes at 4°C to block binding to Fc receptors. Extracellular antigens were stained for 30 minutes at RT in staining buffer. For intracellular protein analysis, cells were fixed and permeabilized with Cytofix (BD Biosciences) and/or Ebioscience Foxp3 kit, per manufacturer instructions. Intracellular antigens were stained for 30min at 4°C in the respective 1x Perm/Wash buffer or overnight for intracellular Aire staining. Cells were analyzed on a Cytek Aurora™.

## Data and code availability

The scRNA-seq datasets are publicly available through the Gene Expression Omnibus database under accession GSE200148, GSE176282, GSE184175, and GSE174405. The code used for downstream analysis and figure generation is accessible at https://github.com/pty0111/TC-data-integration.git.

